# RNA-Seq Analysis Illuminates the Early Stages of *Plasmodium* Liver Infection

**DOI:** 10.1101/870030

**Authors:** Maria Toro-Moreno, Kayla Sylvester, Tamanna Srivastava, Dora Posfai, Emily R. Derbyshire

## Abstract

The apicomplexan parasites *Plasmodium* spp. are the causative agents of malaria, a disease that poses a significant global health burden. *Plasmodium* spp. initiate infection of the human host by transforming and replicating within hepatocytes. This liver stage (LS) is poorly understood when compared to other *Plasmodium* life stages, which has hindered our ability to target these parasites for disease prevention. We conducted an extensive RNA-seq analysis throughout the *Plasmodium berghei* LS, covering as early as 2 hours post infection (hpi) and extending to 48 hpi. Our data revealed that hundreds of genes are differentially expressed at 2 hpi, and that multiple genes shown to be important for later infection are upregulated as early as 12 hpi. Using hierarchical clustering along with co-expression analysis, we identified clusters functionally enriched for important liver-stage processes such as interactions with the host cell and redox homeostasis. Furthermore, some of these clusters were highly correlated to the expression of ApiAP2 transcription factors, while showing enrichment of mostly uncharacterized DNA binding motifs. This finding presents potential LS targets for these transcription factors, while also hinting at alternative uncharacterized DNA binding motifs and transcription factors during this stage. Our work presents a window into the previously undescribed transcriptome of *Plasmodium* upon host hepatocyte infection to enable a comprehensive view of the parasite’s LS. These findings also provide a blueprint for future studies that extend hypotheses concerning LS gene function in *P. berghei* to human-infective *Plasmodium* parasites.

**IMPORTANCE:** The LS of *Plasmodium* infection is an asymptomatic yet necessary stage for producing blood-infective parasites, the causative agents of malaria. Blocking the liver stage of the life cycle can prevent clinical malaria, but relatively less is known about the parasite’s biology at this stage. Using the rodent model *P. berghei*, we investigated whole-transcriptome changes occurring as early as 2 hpi of hepatocytes. The transcriptional profiles of early time points (2, 4, 12, and 18 hpi) have not been accessible before due to the technical challenges associated with liver-stage infections. Our data now provides insights into these early parasite fluxes that may facilitate establishment of infection, transformation and replication in the liver.

## INTRODUCTION

*Plasmodium* spp., the causative agents of malaria, are eukaryotic parasites with a largely conserved and complex life cycle that begins in the mammalian host by invasion of hepatocytes. In these host cells, a single parasite, termed a sporozoite, will transform and then replicate asexually to form thousands of merozoites, or blood-infective forms (1). After maturation and release from the liver, parasites replicate within erythrocytes causing the clinical manifestation of malaria. Some parasites differentiate into sexual forms (gametocytes) that are ingested by an *Anopheles* mosquito during a blood meal. In the mosquito, female and male gametocytes undergo sexual reproduction, and a series of developmental changes lead to a transformation into sporozoites. Inoculation of these sporozoites in the host via a mosquito bite perpetuates the life cycle (2). Despite the significant global burden of malaria (3), our molecular understanding of the *Plasmodium* life cycle is incomplete, hindering our ability to target these parasites to prevent disease and reduce transmission. In particular, the changes that enable sporozoites to transform and then develop within hepatocytes are largely unknown.

Transcriptomic studies have been instrumental in revealing gene expression variation that accompanies stage transitions and developmental processes in *Plasmodium*. Subsequent analyses of these data have also identified transcription factors that are critical for controlling parasite progression at various stages (reviewed in (4)). Yet, only a handful of transcriptome analyses have been completed in the LS relative to other parasite forms, likely owing to the technical challenges associated with studying this stage. Still, these studies have provided important insight into LS-specific biological processes (5), including hypnozoite markers (6, 7), through comparative gene expression analysis with other stages (8), even at single-cell resolution (9). These studies examined gene expression upon the establishment of a LS-trophozoite (24 hours post-infection and thereafter); however, the early stages of LS infection (0–24 hours post-infection) for any *Plasmodium* species remains unresolved.

Our current understanding of the early stages of LS development comes from ultrastructural (10) and immunofluorescence (11) studies. Upon traversal and invasion of hepatocytes, rod-shaped sporozoites expulse unnecessary organelles into the parasitophorous vacuole (PV), which is accompanied by the formation of a protrusion, a bulbous expansion and a transformation into a spherical, replication-competent trophozoite (10). Although this metamorphosis is obvious at the cellular level, the molecular events underpinning this sequence of events remain obscure. Previous studies have examined the gene expression of sporozoites grown axenically since sporozoites can complete this transformation extracellularly if activated by BSA, calcium, and a temperature shift (12, 13). Yet, axenically-grown sporozoites show reduced viability and poor developmental capacity compared to intracellular parasites, suggesting an important role of host pathways in this process. Indeed, a recent study showed that activation of the host GPCR CXCR-4 is necessary for proper parasite metamorphosis (11), highlighting the need to study parasite transformation, and all its subsequent development, in the context of the host cell.

Here, we present a transcriptomic survey of the early and mid-liver stages of *P. berghei* infecting human hepatoma cells. The rodent *P. berghei* and *P. yoelii* LS models are routinely used to study this stage due to their genetic accessibility and tractability relative to human-infective counterparts. Our dataset includes seven timepoints, from 2 to 48 hours post-infection (hpi), making it the most comprehensive transcriptomic analysis of the *Plasmodium* LS to date. We describe changes in gene expression associated with the early stages of *Plasmodium* intracellular development in the LS and show that upregulation of most genes important for exo-erythrocytic form (EEF) maturation occurs as early as 12 hpi. This finding suggests genes important for late-LS development are subject to dynamic expression or translational repression until protein expression is necessary. Furthermore, using co-expression analysis we identified functionally enriched gene clusters with distinct expression patterns and discovered dozens of potential regulatory DNA motifs associated with these genes. Overall, our work completes the life cycle of this important model organism, *P. berghei*, from the transcriptomic perspective, providing a resource for exploring stage-specific expression of genes, and thus providing insight into *Plasmodium* biology.

## RESULTS

### RNA-Seq of Early- and Mid-*P. berghei* Liver Stages

During the course of the LS, sporozoites undergo morphological changes and rapid replication. To investigate differentially expressed transcripts that flux during this stage, HuH7 or HepG2 hepatoma cells were infected with GFP-expressing *P. berghei* ANKA sporozoites. At various times post-infection, samples were harvested, and 1000–3000 *P. berghei*-infected cells were collected by FACS (**Figure 1A**). A poor understanding exists for the early- and mid-LS, therefore greater sampling was acquired before 24 hpi at 2, 4, 12 and 18 hpi (early). Previously analyzed mid-LS samples at 24 and 48 hpi were collected to enable comparison to other studies, as well as 36 hpi, which has not been previously evaluated. *Plasmodium* infection in liver cells is highly heterogenous, with ~ 50% of sporozoites that invade liver cells failing to establish productive infections (14, 15). We ensured selection of populations enriched for productive infections within viable host cells by isolating cells that are both infected and have an uncompromised membrane (GFP^+^ Sytox Blue^-^). FACS analysis indicates that the population of infected cells (GFP-positive) shifts as a function of time, consistent with proper intrahepatic parasite maturation (**Figure 1B**). Further, our gating excluded unviable host cells (Sytox Blue-postive). In our method, we sorted directly into lysis buffer. RNA was then extracted in each sample using the Clontech kit for ultra-low input RNA. Samples were evaluated for concentration and quality using a Qubit and Bioanalyzer, respectively, and analyzed by RNA-seq if they met quality controls. To facilitate robust analysis, sample collection continued until a minimal of 3 replicates per time point were acquired, which yielded a final range of 3–8 replicates.

**Figure 1.**
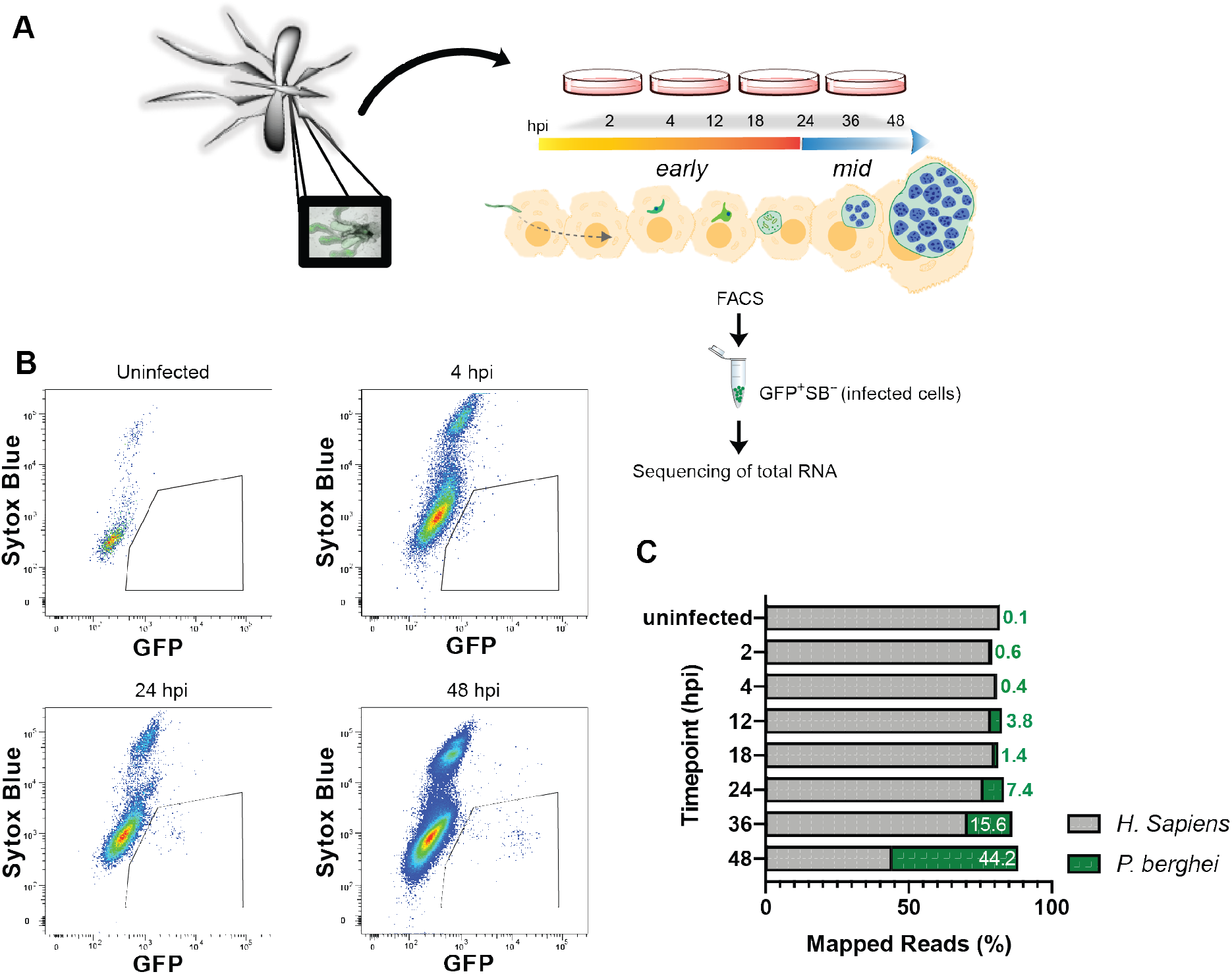
Experimental design for RNA-seq of early and mid-stages of *P. berghei* liver infection. **(A)** Experimental design schematic. Female *Anopheles* mosquitoes were dissected and GFP-expressing *P. berghei* sporozoites were harvested to infect HuH7 or HepG2 cells. Cells were harvested 2, 4, 12, 18, 24, 36 or 48 hpi and FACS-sorted to enrich viable *P. berghei-infected* cells for RNA collection. **(B)** Representative flow cytometry fluorescence dot plots indicating the population of GFP^+^SytoxBlue^-^ cells that were collected at various time points. **(C)** Relative percentage of transcripts mapping to *P. berghei* or *H. sapiens* at various times post infection. Uninfected samples correspond to naïve uninfected cells treated with debris from dissected male *Anopheles* mosquito salivary glands. Data are median of 2–5 biological replicates.

All samples were aligned to *H. sapiens* and *P. berghei* for analysis. As parasite nuclear division does not occur until mid-LS, <4% of the reads mapped to *P. berghei* before 24 hpi. This percentage rises continuously during mid-LS, when the parasite undergoes nuclear division and by 48 hpi ~45 % of the reads correspond to *P. berghei* (**Figure 1C**). In this report, we are focused on parasite processes that control development within hepatocytes, thus principal component analysis (PCA) was completed on *P. berghei* data after removal of batch effects. PCA revealed no major differences between the parasite transcriptomes obtained by infecting HepG2 or Huh7 cells (**Figure S1A**), but a general clustering of replicates by genotype (timepoint) was observed (**Figure S1B**). Of note, PCA showed strong separation of 4 hpi and 2 hpi, with the latter grouping well with sporozoites, highlighting the parasite transformations that must occur during these 2 hours.

To analyze our dataset in the context of the entire *Plasmodium* life cycle, we calculated Spearman correlations on our data as well as previously published *Plasmodium* transcriptomic data from sporozoites, the asexual blood stage (ABS), gametocytes, ookinetes, hypnozoites and the LS (**Table S1**). This analysis spanned data obtained from *P. berghei, P. yoelii, P. cynomolgi, P. vivax* and *P. falciparum*. Consistent with previous reports, the LS was more similar to the asexual blood stage (ABS) than to gametocytes and ookinetes (8). Indeed, we observe two general groups comprising of 1) mostly metabolically active, intracellular stages (LS, ABS), and 2) mostly motile, extracellular stages (**Figure 2**). Notably, early liver stages of *P. berghei* and (axenic) *P. vivax* (LS_2h/4h) fell into the latter group, being more highly correlated to sporozoites, ookinetes and gametocytes than to other LS time points.

**Figure 2.**
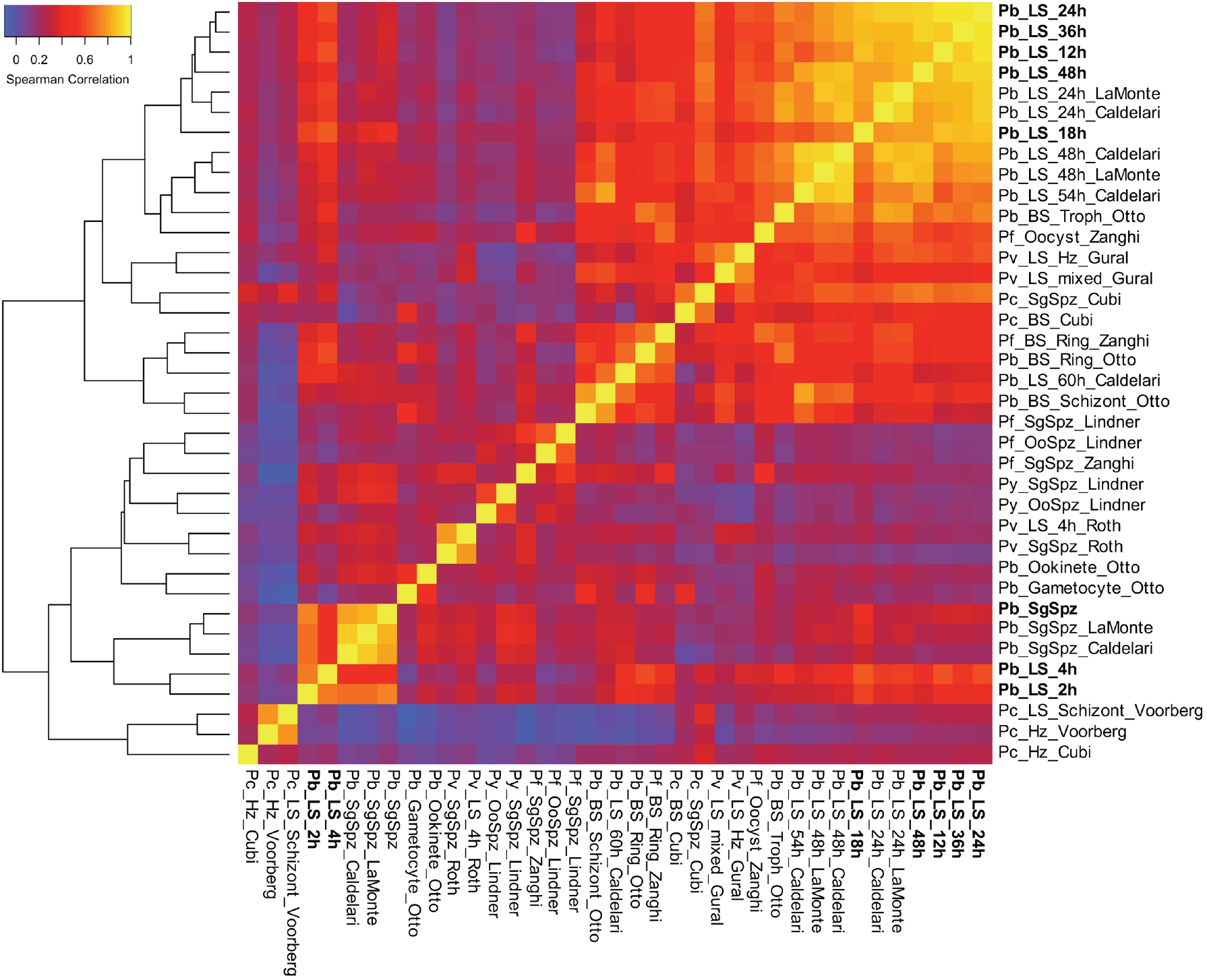
Overview of *Plasmodium* transcriptome analyses. Hierarchical clustering of gene expression datasets from different stages of the *Plasmodium* life cycle (7, 8, 26, 32, 34, 52–54). Datasets generated in this study are in bold. Clustering is based on Spearman correlation coefficients calculated and plotted using R. Refer to Table S1 for information regarding the datasets used to generate this figure.

### Early Liver-Stage Transcriptome of *P. berghei*

Thousands of statistically significant differentially expressed transcripts were detected at early LS time points, with most of these transcripts being downregulated at 2 and 4 hpi, and then upregulated at 12 hpi with respect to sporozoites (**Figure 3A and B, Data S1**). This shift suggests a change from gene suppression to activation as the parasite exits the early stage of intrahepatic development. As expected, genes important for host cell traversal and invasion, such as *CELTOS, SUB2*, and *CSP*, were downregulated at 2 hpi, concurrent with upregulation of genes important for nutrient acquisition (*ZIP1, TPT, NT1*), reflecting the establishment of the infection in the host cell. Unsurprisingly, at these early stages, we also observe strong upregulation of *EXP2* and *PV2*, which encode PVM-associated proteins, together with several predicted exported proteins with unknown function, indicating that early (<4 hpi) establishment and remodeling of the PVM is essential for parasite LS maturation. Interestingly, we observe that *LYTB* (IspH), the last enzyme in the isoprenoid biosynthesis pathway in the apicoplast, is among the most upregulated genes at both 2 and 4 hpi (**Table S2**). Apicoplast pathways are important potential drug targets for the development of LS antimalarials, but are not known to be involved in early-LS processes.

**Figure 3.**
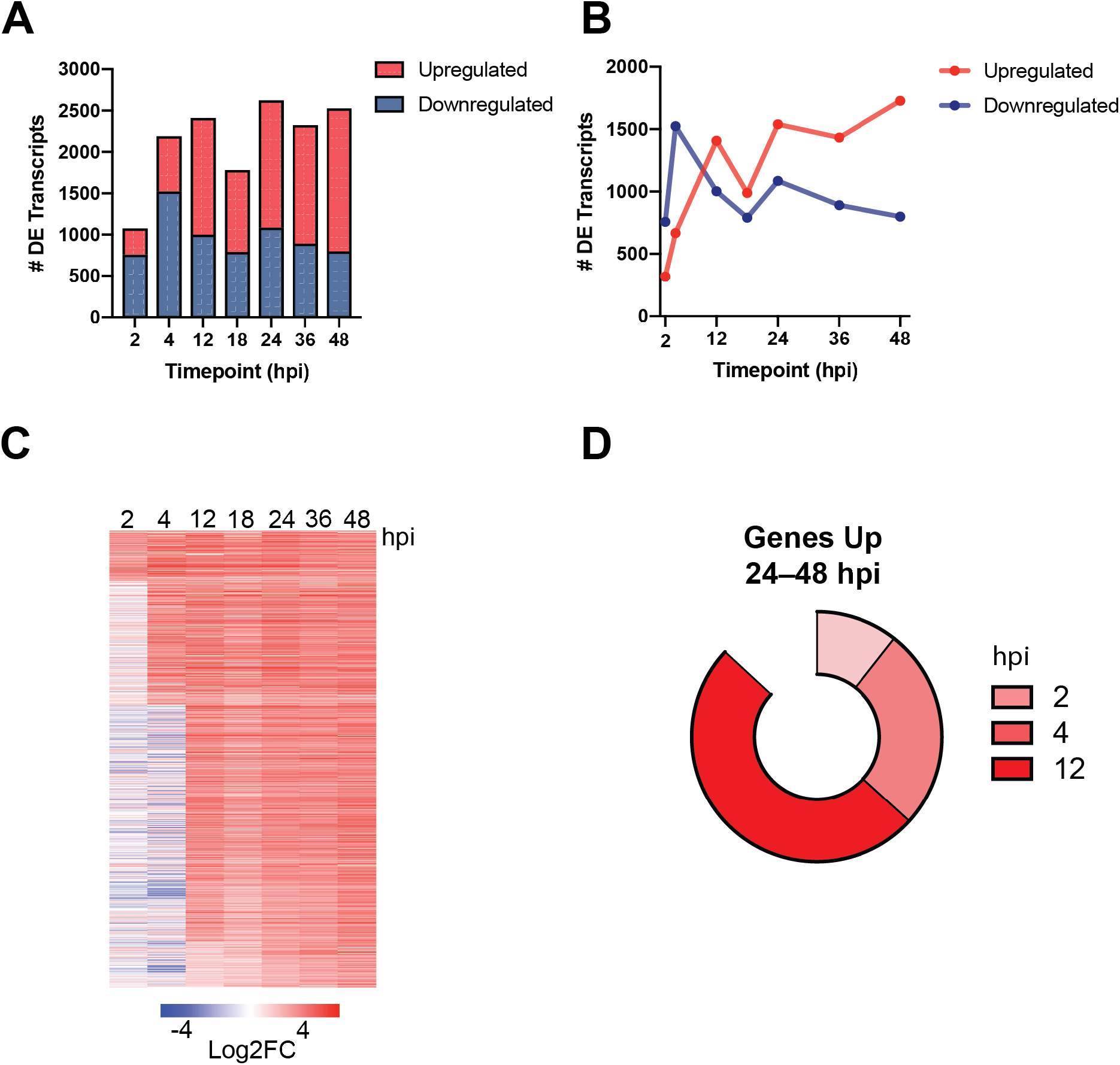
Dynamic gene regulation throughout liver-stage *P. berghei* development. Total **(A)** and upregulated (red)/ downregulated (blue) **(B)** differentially expressed (DE) transcripts (*q* < 0.01) shown at each time point. **(C)** Expression profiles of 1,197 genes upregulated at 24, 36 and 48 hpi ordered based on the timepoint they were first observed to be upregulated. Expression is shown as the Log2 fold change vs sporozoite samples. **(D)** The proportion of genes that are upregulated throughout late stage development (24, 36 and 48 hpi) are divided by when they are first observed to be upregulated (2, 4 or 12 hpi).

Translational regulation of *Plasmodium* transcripts has been extensively documented and it is known to play a pivotal role during developmental transitions in the life cycle. We found pervasive upregulation of most of the functionally characterized translational regulators in *Plasmodium*, at the exclusion of *PUF1* and *PUF2*, which appeared to be dramatically downregulated when compared to their high expression in sporozoites*. DOZI, ALBA1, ALBA2* and *ALBA4* were upregulated as early as 4 hpi (Log2FC < 2, *q* < 0.01) (**Figure S2**). Moreover, among the highest differentially expressed transcripts at 2 and 4 hpi, there was an enrichment of genes involved in RNA-protein complexes and interactions, such as *SR1, NOP10, CBF5, RPS12, NAPL* (**Table S3**). Thus, translational regulation likely plays an important role in the early stages of *Plasmodium* infection of the liver.

At ~24 hpi and thereafter, the single-nucleated trophozoites replicate and subsequently mature into LS schizonts, each harboring tens of thousands of nuclei. Previous work examining the LS stage transcriptome at these mid-stages identified hundreds of differentially expressed genes involved in translation, metabolism, protein trafficking and redox processes (5, 8). Since we saw a strong correlation between 12 hpi and mid-liver stages (Spearman correlation = 0.837— 0.949, **Figure 2**), we questioned how early a statistically significant upregulation of the core mid-LS transcriptome could be observed in our dataset. We found 1,197 genes in our dataset that are significantly upregulated at 24, 36 and 48 hpi compared to sporozoites (*q* < 0.01), constituting about 20% of the *P. berghei* genome (**Figure 3A** and **B**). Interestingly, we find that 87% of transcripts that are upregulated throughout the mid-liver stages (24 hpi through 48 hpi) are upregulated as early as 12 hpi (**Figure 3C**). More specifically, 50% of the genes that are upregulated in the mid-liver stages are first observed to be upregulated at 12 hpi (**Figure 3D**).

### Co-Expression Analysis Identifies Functionally Enriched Gene Clusters

To identify coexpression patterns that may inform future functional studies, we performed a clustering analysis of the k-means for all differentially expressed genes for all of the samples included in our dataset. Fourteen clusters emerged from this hierarchical clustering analysis (**Figure 4A, Data S2**). These clusters could be further grouped within three major co-expression patterns when columns were grouped by sample genotype (timepoint). The first major cluster group (clusters 3, 11 and 13) includes genes that are upregulated early during infection (spz, 2 and 4 hpi) and are generally downregulated throughout the rest of LS infection, such as *ETRAMPs* and *SPELD*. The second major cluster group (clusters 1, 2, 4, 7, 8, 9, 12, and 14) includes genes that are downregulated during the early stages of infection, but are then consistently upregulated from 24 to 48 hpi. The third major cluster group (clusters 5, 6, and 10) includes genes that are upregulated throughout the entire LS.

**Figure 4.**
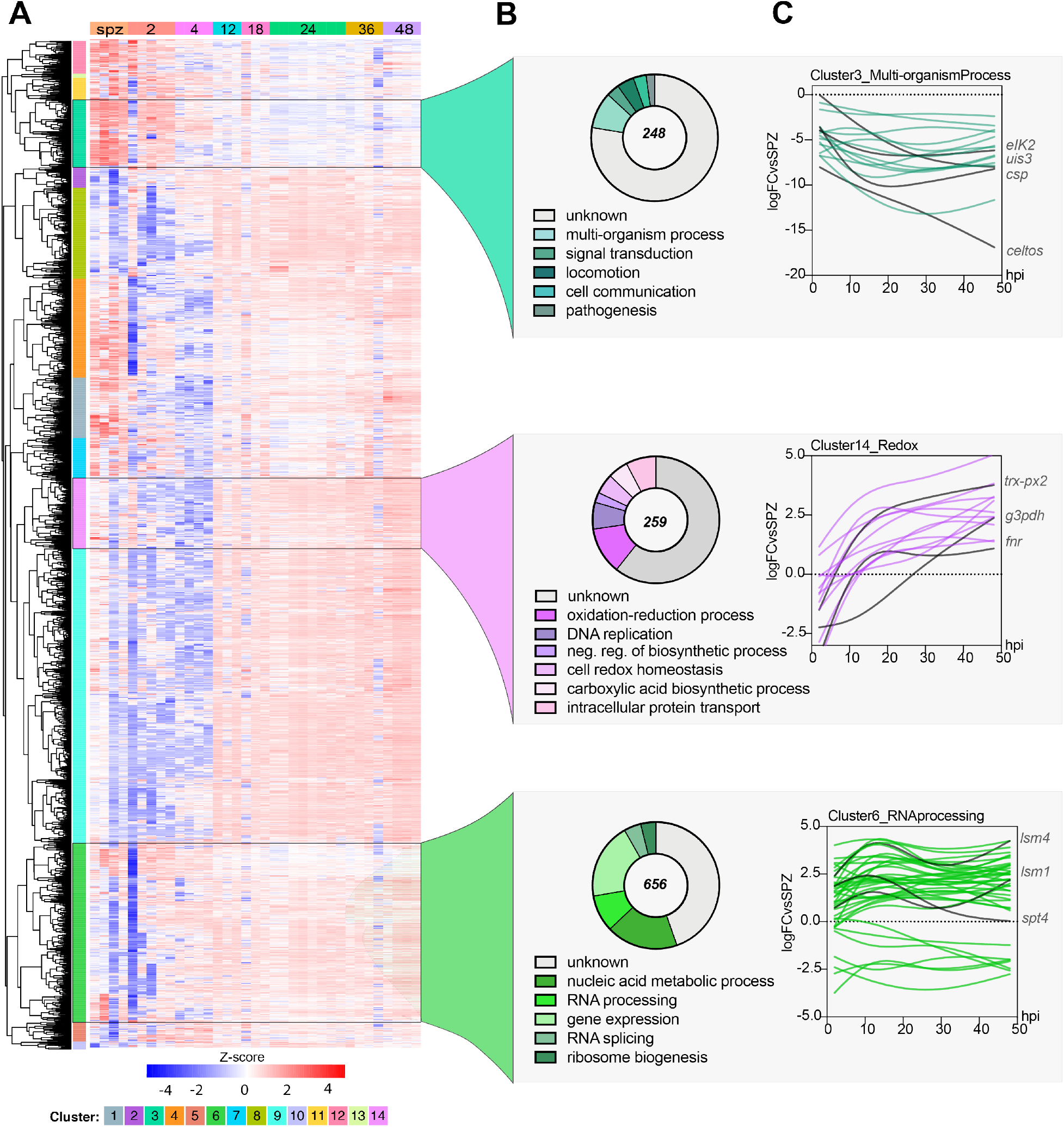
Co-expression analysis identifies enriched processes during *P. berghei* development in hepatocytes. **(A)** Hierarchical clustering using a correlation distance with complete linkage of all genes significant (FDR <= 5%) in at least one of the analyses. Gene expression is z-score transformed. **(B)** GO Enrichment analysis (biological process) of enriched clusters 3, 14 and 6 shown. Representative GO terms (*p* < 0.01) and their respective number of genes (pie chart) are shown. Total number of genes in each cluster is shown at the center of the pie chart. **(C)** Spline models of gene expression data for all the genes in the top-scoring GO term in each cluster. Key genes in each group and their expression patterns are highlighted in red. Refer to **Data S3** for complete GO analysis of all clusters.

To investigate possible enrichment of biological processes of co-expressed genes, we analyzed each cluster by gene ontology (GO). Such analyses revealed the enrichment of various GO terms for each of the clusters (*p* < 0.01). We prioritized clusters for which at least one GO term was enriched by a *p-adj* (Bonferroni) < 0.01. Cluster 3 stood out as highly enriched despite 142 out of the total 248 genes in this cluster not being annotated. For this cluster, enrichment analysis indicated significant enrichment of “interspecies interaction” (GO:0044419, *p* < 1.91E-07) as well as locomotion (GO:0040011,*p* < 0.0005679) and signal transduction (GO:0007165,*p* < 0.00049572) (**Figure 4B**). Genes in this cluster are highly expressed in sporozoites, and thus appear to be strongly downregulated during infection (**Figure 4C**). In agreement with this result, this cluster includes genes that have been previously shown to play an important role during invasion (*CELTOS, SPECT1, TRAP*), interactions with the host liver cell (*UIS3, UIS4, CSP, p36, p52*), and translational control of liver-stage specific transcripts (*UIS2, PUF1, PUF2*) (16).

Cluster 14 was enriched for “oxidation-reduction process” (GO:0055114, *p* < 9.01E-05), “DNA replication” (GO:0006260, *p* < 0.00164223), and “intracellular protein transport” (GO:0006886, *p* < 0.00941436) (**Figure 4B**). In this group, genes involved in redox-regulatory processes (*FNR, TRX-PX2*), as well as biosynthetic genes such as *G3PDH*, can be found. Expression of genes under the redox group appears to peak by ~12 hpi and then remains stably upregulated throughout infection. This expression pattern highlights the need for this machinery to mitigate potential stress due to the dramatic parasite replication and growth that is initiated at ~24 hpi (**Figure 4C**). Little is known about redox biology in *Plasmodium* parasites, particularly during the LS, but these processes have historically been key pathways for drug discovery. Indeed, atovaquone, a drug for malaria prophylaxis in combination with proguanil, inhibits liver stage parasites in vitro by impairing mitochondrial redox metabolism via targeting the cytochrome *bc1* complex (17). This dataset may serve as a starting point to discover more LS targets involved in redox metabolism. Furthermore, although not enriched in our GO analysis, we observed that several important liver-specific genes are found in this cluster, such as *IBIS1*, *LISP1*, and *LISP2*. Finally, in cluster 6, we saw enrichment of core functions such as “gene expression” (GO:0010467, *p* < 5.71E-05) and “RNA processing” (GO:0006396,*p* < 5.98E-06), which contains 656 genes. As expected of housekeeping functions, these genes appear to be expressed throughout LS infection.

Our analysis identified several clusters with enriched GO terms, some which accurately describe the known LS biology at different timepoints. Although GO enrichment provided a useful assessment of differentially expressed processes, we note that it is limited in its reach in *Plasmodium* when compared to other model organisms since ~40% of the genome remains unannotated. Hence, to further explore the composition of these co-expression clusters, we made use of the Rodent Malaria genetically modified parasite database (RMgmDB) to provide phenotypic information about our clusters throughout the life cycle (18). Interestingly, we observe that while most clusters have a high proportion of genes for which disruption resulted in phenotypes across the entire life cycle, only a few clusters had genes that displayed phenotypes exclusively in sporozoite and/or liver stage (**Figure S3**). Specifically, clusters 3 and 14 had the highest percentage of spz/LS-specific genes (13 and 9%, respectively), reinforcing the potential for identifying new LS drug and vaccine targets within these clusters.

### Expression Dynamics of AP2 Transcription Factors

Transcriptional regulation of gene expression has been extensively studied in the intraerythrocytic developmental cycle (IDC) and mosquito stages of *P. berghei* and *P. falciparum*. The AP2 transcription factors (TFs), comprised of 26 genes in *P. berghei*, are the best-characterized family of TFs in apicomplexans (**Figure 5A**). AP2s are known to regulate *Plasmodium* transitions into different developmental stages and have emerged as key factors leading to both sexual commitment and sex differentiation (reviewed in (4)). Unsurprisingly, we observe that AP2 genes with established functions in mosquito stages (*AP2-O*, *AP2-O2*) and those involved in sporozoite development (*AP2-SPs*) are downregulated throughout the liver stages (**Figure 5A**). The only ApiAP2 TF known to play a role in LS development is *AP2-L*. *AP2-L* (-/-) parasites are able to traverse and invade liver cells, but arrest in the schizont stage (19). *AP2-L* transcripts are abundant in sporozoites and thus appear to be strongly downregulated during infection, as early as 2 hpi (**Figure 5A**).

**Figure 5.**
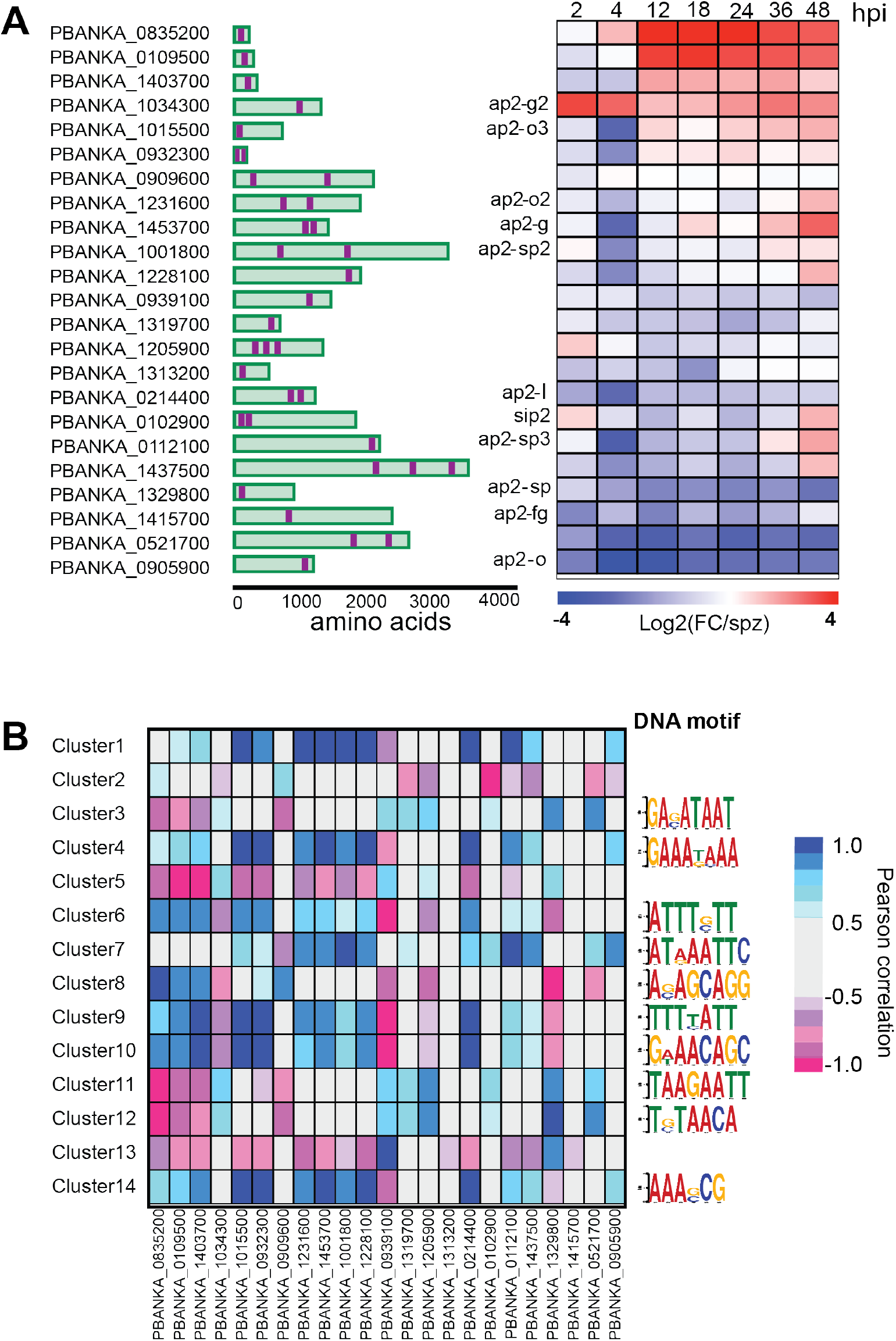
Expression of *P. berghei* AP2 transcription factors in the liver stage. **(A)** Gene IDs of the 26 AP2 transcription factors in the *Pb* genome, their respective protein architecture schematic (with AP2 displayed in purple) and their corresponding expression as the log2 fold change vs spz at each time point in the LS. **(B)** Heatmap of Pearson correlations between AP2 transcription factors and the average expression of all genes in each cluster (left). The top most enriched DNA motif for each cluster discovered through the DREME pipeline is shown (right). Refer to **Data S3** for the complete set of motifs and their respective enrichment score.

We observe strong upregulation (~3-fold) of *AP2-G2* at 2 and 4 hpi. AP2-G2 has been shown to act as a repressor during the BS and gametocyte development, and to have different targets during these stages (20, 21). A group of such targets corresponds to the liver-specific genes *LISP1* and *TREP*, which are important for LS schizont maturation and expressed during late LS infection (21). Interestingly, we observe that *AP2-G2* expression is negatively correlated to the average expression of the main clusters harboring this set of genes, including clusters 1, 9 and 14 (**Data S3**). Thus, it is plausible that during the first hours of infection, *AP2-G2* acts as a repressor of genes involved in later stages of LS development, many of which remain uncharacterized.

Interestingly, we observed significant upregulation of the uncharacterized ApiAP2s PBANKA_0835200 and PBANKA_0109500 throughout the LS starting at 12 hpi, in contrast to the early upregulation of *AP2-G2*. While still functionally uncharacterized, their orthologs in *P. falciparum* have been recently shown to co-express during differentiation in gametocytogenesis, and to be inversely correlated to genes involved in ABS development. This expression pattern suggests they may have a role as co-repressors of genes involved in the ABS (22). In our dataset, we observe a strong correlation with clusters 11, 12 (both negative) and cluster 8 (positive).

We sought to identify enriched DNA motifs in each of the co-expression clusters by analyzing the 5’ UTR sequences (1 kb) of their genes against the upstream sequence for all of the genes in other clusters using DREME (**Data S3**) (23). While genes in clusters 1, 2, 5 and 13 lacked any enriched DNA motifs, *de novo* discovery uncovered hundreds of DNA motifs in the remaining clusters, with the topmost enriched motif shown in **Figure 5B**. We found that the most significant motif in cluster 12 (T[G/C]TAACA) matched the motif recognized by ApiAP2 PBANKA_0521700 (GTGTTACAC, *p* < 1.28e-05). This cluster included genes that are mostly downregulated throughout the LS until the later time points in our time series, such as the BS schizont-specific genes *SERA2* and *SERA3*. Additionally, PBANKA_0521700 expression was strongly correlated to cluster 12 (r = 0.83, *p* < 0.021), suggesting this cluster might harbor previously unknown targets of this ApiAP2 (**Figure 5B**).

## DISCUSSION

Our data provide novel insights into gene expression fluxes throughout *Plasmodium* development within hepatocytes. The transcriptional blueprints provided by our time series enables comparison of early-, mid- and late-liver stage parasite processes for the first time. We found 146 genes exclusively upregulated early, such as *EIF5;* and 482 genes, including *SERA1* and *LISP2*, exclusively upregulated in the mid-liver stages (**Figure S4**). Furthermore, our datasets recapitulated well-established gene expression patterns of key LS genes, and overall were largely in agreement with recently reported datasets, supporting the validity of our approach. Through our analysis, we identified a key shift in parasite gene expression that occurs at 12 hpi, and the role of transcription factors in driving LS maturation. Specifically, we explored potential transcriptional regulation of co-expressing genes by analyzing their upstream sequences for enrichment of potential DNA binding motifs, and their correlation to *P. berghei* AP2 transcription factors. Our results revealed an association between the uncharacterized PBANKA_0521700 AP2 TF and cluster 12. PBANKA_0521700 is preferentially expressed in the ring stages of the IDC and is refractory to disruption in the BS (24, 25), hampering functional studies of this gene. Our data, in conjunction with previously reported *P. berghei* RNA-seq (8, 24–26) and single cell studies covering the entire life cycle (9), could be useful to refine hypotheses about the functions and targets of this TF as well as other AP2 TFs.

While AP2 TFs have been at the center of gene expression studies in *Plasmodium*, novel “omics” approaches have begun uncovering other layers of gene regulation. Indeed, post-transcriptional regulation, such as *N*^6^-methyladenosine (m^6^A) of mRNA and alternative splicing, have recently been recognized as essential for fine-tuning gene expression in blood and sexual stages (27, 28). In particular, disruption of the splicing factor *Pb*SR-MG was shown to perturb sexspecific alternative splicing, thus demonstrating its role as a cellular differentiation regulator (29). Interestingly, we observed a dramatic upregulation of the splicing factor SR1 coinciding with the parasite’s metamorphoses in the LS, hinting at an important role for alternative splicing during this stage. Future reverse genetic studies may help establish a role for alternative splicing in the LS.

A well-documented form of gene expression regulation in *Plasmodium* occurs at the translational level. Translational repression (TR) of hundreds of transcripts has been reported at most stages of the *P. berghei* life cycle (30). TR is particularly pervasive in the sporozoite transition from the mosquito to the mammalian host (31, 32). During this transition, hundreds of transcripts that are highly expressed in sporozoites are stored in mRNA granules, until infection of the host relieves this repression resulting in protein translation. The extent to which a TR program operates in the LS is currently unknown. However, we observed that ~50% of all transcripts upregulated after 24 hpi are also upregulated at 12 hpi, including some with known roles in LS schizont maturation (*IBIS1* and *BP2*). Furthermore, we see upregulation of several known translation regulators *DOZI, ALBA1,-2,-4*, which could potentially repress translation of transcripts important for late LS development and/or the subsequent transition to the ABS. Unfortunately, this possibility will be exceedingly difficult to test in the absence of robust global proteomic analysis of the early LS parasite. Nonetheless, our data, coupled with recent RNA-seq and proteomic studies of the more accessible late LS, can provide a starting point to address this question (8, 33).

Previous work examining the transcriptional changes of axenically grown early LS *P. vivax* identified upregulation of calcium-related proteins (RACK1) and RNA-binding proteins (*ALBA1, −2* and *−4*) (34). We saw upregulation of the *P. berghei* orthologs of these genes as well as hundreds of other genes, dramatically expanding the dataset for genes upregulated at this stage (**Figure S5**). For example, we found *LYTB* is upregulated at 2 and 4 hpi, indicating isoprenoids may be important at this time. While the FASII and *de novo* heme biosynthesis pathways have been genetically and chemically validated as essential to the late liver stages, less is known about isoprenoid biosynthesis during the early liver stages (35, 36). When intracellular sporozoites metamorphosize to replication-competent trophozoites, most organelles are expelled at the exclusion of the nucleus, mitochondrion, and apicoplast (10). Thus, it is plausible that the apicoplast serves an important metabolic role with isoprenoids in the liver stages of infection. Unfortunately, the use of isoprenoid biosynthesis inhibitors has yielded inconclusive results about its function during the LS (37, 38), emphasizing the need for future genetic studies to elucidate the role of isoprenoid biosynthesis throughout intrahepatic development. Thus, we anticipate our data will be useful to guide future reverse genetic and functional studies to investigate the role of *Plasmodium* genes with important early- and mid-LS functions.

Our understanding of *Plasmodium* LS biology still lags behind that of other parasite life cycle stages, hindering the development of much-needed prophylactic measures to combat malaria. Our work represents a window into the previously undescribed transcriptome of the early LS upon host cell infection and offers a comprehensive view of the *Plasmodium* LS. Future studies expanding on our analysis and validating time-specific LS genes will further advance our molecular understanding of this critical step in the *Plasmodium* life cycle.

## MATERIALS AND METHODS

### Parasites

Sporozoites were freshly harvested prior to experiments from dissected salivary glands of *Anopheles stephensi* mosquitoes infected with *P. berghei* ANKA stably expressing a green fluorescent protein (GFP) purchased from the New York University Langone Medical Center Insectary.

### Cell culture

HepG2 were purchased from ATCC and HuH7 cells were a kind gift from Dr. Peter Sorger (Harvard Medical School). Hepatocytes used for *P*. *berghei* infections were maintained in Dulbecco’s Modified Eagle Medium (DMEM) with L-glutamine (Gibco) supplemented with 10% heat-inactivated fetal bovine serum (HI-FBS) (v/v) (Sigma-Aldrich) and 1% antibiotic-antimycotic (Thermo Fisher Scientific) in a standard tissue culture incubator (37°C, 5% CO_2_).

### Sample collection for RNA-seq

Infected hepatoma cells were collected as previously described (39). Briefly, T25 flasks were seeded with 3×10^5^ HepG2 or 8×10^4^ HuH7 cells. About 24 hours after seeding, cells were infected with 1×10^5^ GFP-expressing *P*. *berghei*-ANKA sporozoites. Infected cells and uninfected controls were sorted directly into RNA lysis buffer (Clontech) using the BD FACSAria II cell sorter (BD Biosciences) at the Duke Human Vaccine Institute. Sytox blue was used as a live/dead cell indicator (Thermo Fisher Scientific). Infected cells were collected by sorting of the GFP, and gated compared to uninfected hepatoma cells. RNA was extracted using SMART-seq v4 Ultra Low Input RNA Kit for Sequencing (Clonetech) and libraries were prepared at the Duke Next Generation Sequencing Core Facility and sequenced on the Illumina HiSeq 4000 as 50 base pair single-end reads. Samples (4–5) were pooled on each flow cell lane.

### RNA-seq and differential expression analysis

RNA-seq data were processed using the TrimGalore toolkit (40) which employs Cutadapt (41) to trim low-quality bases and Illumina sequencing adapters from the 3’ end of the reads. Only reads that were 20 nt or longer after trimming were kept for further analysis. Reads were mapped to a combination of the GRCh37v75 (42) version of the human genome and the PbANKAv3 of the *P. berghei* genome using the STAR RNA-seq alignment tool (43). Reads were kept for subsequent analysis if they mapped to a single genomic location. All samples mapping >1 million reads to the *P. berghei* genome were used for a preliminary analysis. Gene counts were compiled using the HTSeq tool (44). Only *P. berghei* genes that had at least 10 reads in any given library were used in subsequent analysis. Normalization and differential expression were carried out using the DESeq2 (45) Bioconductor (46) package with the R statistical programming environment (47). The false discovery rate was calculated to control for multiple hypothesis testing. When calculating the differential expression between genes at each time point relative to the control, the cell type and sequencing batch were included as cofactors in the model.

Spearman correlations between published *P. berghei* RNA-seq datasets and our own were calculated and plotted using the *cor* function in the *stats* R package.

### Clustering analysis

To determine the different patterns of gene expression across all groups of samples, we first identified genes that showed differential expression in at least one of the comparisons performed (FDR <= 5%). The genes were the clustered across all samples by a correlation distance using complete linkage after z-score transformation. The *NbClust* (48) package was used to separate the gene expression across all samples into distinct clusters.

De novo motif discovery was performed using DREME from the MEME suite (23). For each cluster the input data set was the upstream 1000 kb region of each gene within that cluster, and the negative set was the upstream region of genes that were not in that cluster. The analysis was run in discriminative mode, scanning the given strand only, with the predicted motif size of 4–10 bp and cut-off E value of 0.05. The top most enriched motif for each cluster was then analyzed with TOMTOM (49) to compare to previously *in silico* discovered motifs (50).

The correlation matrix was generated in PRISM by calculating the Pearson correlation between each AP2 transcription factor and the average fold change expression for all the genes in each cluster.

### Gene Ontology (GO)

GO analyses for each cluster were performed using the GO enrichment tool of Biological Processes in PlasmoDB (51) with a cutoff of *p* <0.01. The number of genes from the cluster in each of the representative top-scoring GO terms (lowest *p*-values) were plotted.

## Supporting information

Dataset 1. Differential expression

Dataset 2. Cluster assignment

Dataset 3. Cluster analysis

## ACKNOWLEDGEMENTS

This work is funded by the NIH (DP2AI138239, to E.R.D), the CM Hauser Fellowship (M.T.M), and the NSF (DGE-1644868, to K.S.). The content of this study is solely the responsibility of the authors and does not necessarily represent the official views of the NIH.

We thank Prof. Ana Rodriguez and Sandra Gonzalez from the NYU Insectary for providing *Plasmodium*-infected mosquitoes, David Corcoran from the Duke Genomic Analysis and Bioinformatics (GCB) Core Facility, the DHVI Flow Cytometry Core Facility, and Joseph Saelens. We also thank Prof. Photini Sinnis, Amanda Balaban, and the JHMRI Insectary and Parasitology Core Facilities for their help. We thank Luisa Toro Moreno for data managing support, and Profs. Jen-Tsan Ashley Chi and Steven Haase for useful discussions.

## Author contributions

Conception or design of the work— M.T-M, K.S, D.P, E.R.D

Data collection—K.S, D.P, E.R.D

Data analysis and interpretation— M.T-M, K.S, T.S

Drafting the article— M.T-M

Critical revision and contributions to the article— K.S, E.R.D

Final approval of the version to be published— M.T-M, K.S, T.S, D.P, E.R.D

