## Supplementary Information for "RNA-Seq Analysis Illuminates the Early Stages of *Plasmodium* Liver Infection"

### SUPPLEMENTARY MATERIALS

#### Figures and Tables

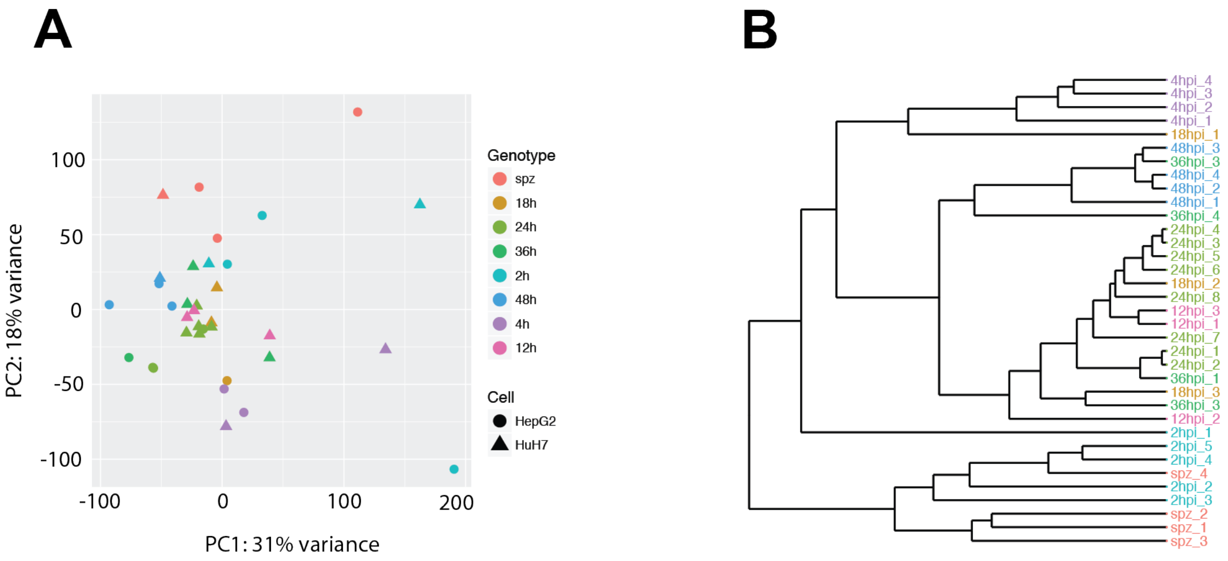


**Figure S1. (A)** PCA of all samples after removal of batch effects. Colors correspond to genotypes/timepoints (spz as well as 2, 4, 12, 24, 36, and 48 hpi). Shapes designate the cell line used for the infection. **(B)** Hierarchical clustering of the different samples based on all genes using a correlation distance with complete linkage. The sample names are colored by genotype (legend in left panel). The PCA showcases major sources of variation in the data, while the hierarchical clustering plot specifically identifies the most similar samples (based on correlation of all genes) and groups them together in a step-wise fashion. For the PCA analysis, we showcase the first two components to illustrate that while the largest source of variation does not separate out the time-points (genotypes), it does show that cell type and sequencing batch are not major contributors to the variation of the data. Nonetheless, the hierarchical clustering shows that, overall, our time-points tend to cluster together, based on correlation of all genes.

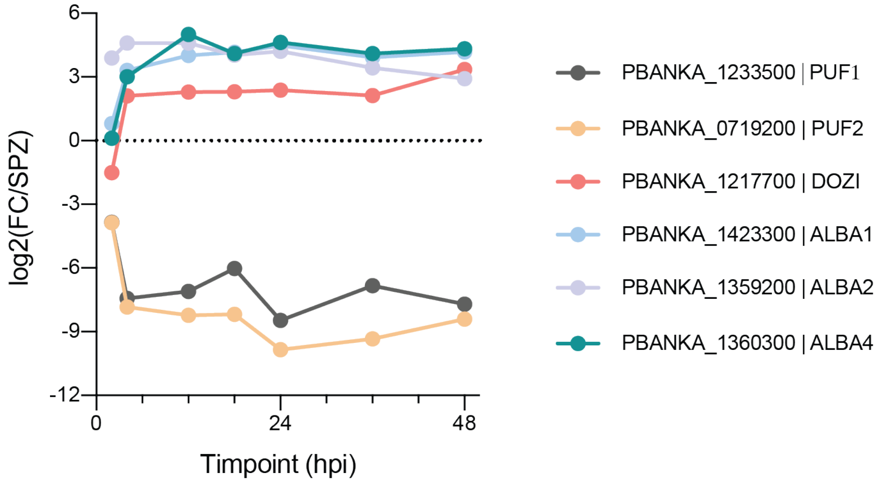


**Figure S2.** Expression profile of *P. berghei* translation regulators in the liver stage. The gene IDs and names are shown in legend. Expression is shown as the Log2(fold-change).

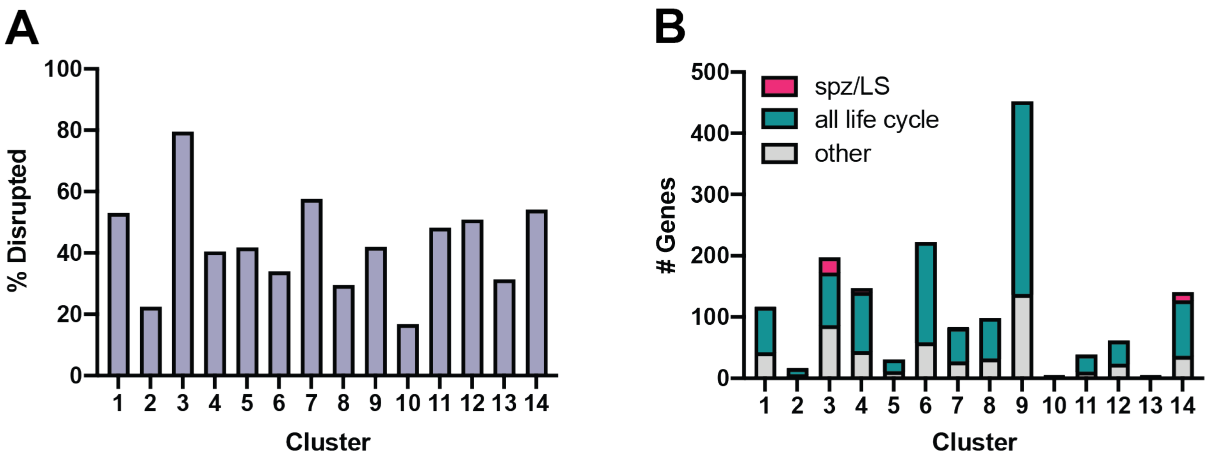


**Figure S3.** Phenotypic overview of co-expression clusters. **(A)** Percent of genes in each cluster that are reported to have been "successfully disrupted" in the RMgmP database. **(B)** Number of genes in each cluster for which successful disruption resulted in distinct phenotypes from WT parasites. "Spz/LS" (pink) denotes mutants displaying phenotypes only in sporozoites or liver stages; "all life cycle" (teal) display phenotypes across all life cycle (asexual blood stage, gametocyte/gamete, fertilization and ookinete, oocyst, sporozoite, and liver stage); “other” (grey) display phenotypes distinct from the two above. All data are from the RMgmDB (18).

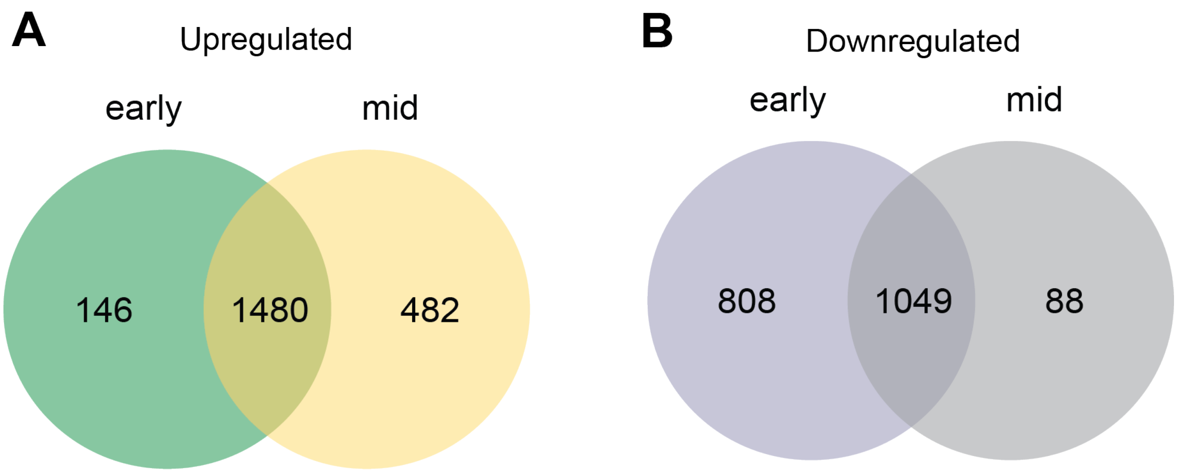


**Figure S4**. Venn diagram illustrating the overlap of the upregulated (**A**) and downregulated (**B**) genes at early (2,4,12, or 18 hpi) and mid-stages (24, 36, or 48 hpi).

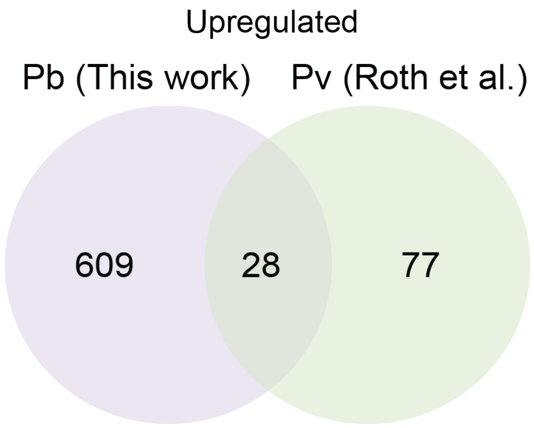


**Figure S5**. Venn diagram illustrating the overlap of upregulated genes from our 4 hpi dataset and Roth *et. al* 's *P. vivax* early liver-stage data (RPMI, 0 h vs. RPMI 4 h, 37 °C). *P. berghei* genes were translated into their respective syntenic *P. vivax* orthologs for this comparison (Hypergeometric *p*-value < 3.4e-06). We speculate that we don't see more overlap in upregulated genes because of 1) technical and experimental issues, such as the differences in coverage between the two datasets and the statistical cutoffs used to determine differential expression and 2) biological differences, such as axenic culture vs intracellular developmental, and perhaps distinct temporal regulation of developmental programs between *P. vivax* and *P. berghei*.

**Table S1.** Datasets used in Spearman correlation analysis (Figure 2).

| Source (ref) | Parasite Species | Samples | Host cell |
|---|---|---|---|
| Zanghi (54) | *P. falciparum* | SgSpz, oocyst, ring | |
| Gural (53) | *P. vivax* | hypnozoite (Hz), mixed LS | Primary human hepatocytes |
| Voorberg (55) | *P. cynomolgi* | LS (schizont), hypnozoite (Hz) | *M. mulatta* or *M. fascicularis* hepatocytes |
| Cubi (7) | *P. cynomolgi* | Hypnozoite (Hz), SgSpz, BS | *M. fascicularis* hepatocytes |
| Caldelari (8) | *P. berghei* | SgSpz, LS_24, LS_48, LS_54, LS_60 | HeLa |
| LaMonte (26) | *P. berghei* | SgSpz, LS_24, LS_48 | HuH7, HC04, HepG2 |
| Lindner (32) | *P. falciparum, P. yoelii* | SgSpz, OoSpz | |
| Roth (34) | *P. vivax* | SgSpz(0h), LS_4h | |
| This work | *P. berghei* | SgSpz, LS_2h, LS_4h, LS_12h _LS_18h, LS_24h, LS_36h, LS_48h | HuH7 and HepG2 |

**Table S2.** Top most upregulated genes at 4 hours post-infection.

| Gene ID | Product Description | Gene Name |
|---|---|---|
| PBANKA_0201500 | Plasmodium exported protein, unknown function | N/A |
| PBANKA_0208700 | 4-hydroxy-3-methylbut-2-enyl diphosphate reductase, putative | LytB |
| PBANKA_0214600 | Plasmodium exported protein, unknown function | N/A |
| PBANKA_0315420 | Plasmodium RNA of unknown function RUF2 | N/A |
| PBANKA_0405400 | 40S ribosomal protein S12, putative | RPS12 |
| PBANKA_0405500 | 60S ribosomal protein L7, putative | N/A |
| PBANKA_0418500 | Plasmodium exported protein, unknown function | N/A |
| PBANKA_0517100 | conserved protein, unknown function | N/A |
| PBANKA_0602700 | nucleosome assembly protein, putative | NAPL |
| PBANKA_0619700 | rhoptry-associated leucine zipper-like protein 1, putative | RALP1 |
| PBANKA_0623200 | lysophospholipase, putative | N/A |
| PBANKA_0623500 | fam-a protein | N/A |
| PBANKA_0700500 | fam-a protein | N/A |
| PBANKA_0807100 | stomatin-like protein | STOML |
| PBANKA_0942100 | metabolite/drug transporter, putative | N/A |
| PBANKA_1011900 | H/ACA ribonucleoprotein complex subunit 3, putative | NOP10 |
| PBANKA_1025200 | H/ACA ribonucleoprotein complex subunit 4, putative | CBF5 |
| PBANKA_1030700 | conserved Plasmodium protein, unknown function | N/A |
| PBANKA_1101100 | Plasmodium exported protein, unknown function | N/A |
| PBANKA_1120800 | signal recognition particle subunit SRP68, putative | SRP68 |
| PBANKA_1127700 | nicotinate phosphoribosyl transferase, putative | NAPRT |
| PBANKA_1302241 | tRNA Selenocysteine | N/A |
| PBANKA_1326400 | glyceraldehyde-3-phosphate dehydrogenase | GAPDH |
| PBANKA_1347800 | 20 kDa chaperonin, putative | CPN20 |
| PBANKA_1365680 | fam-c protein | N/A |
| PBANKA_1407600 | 60S ribosomal protein L24, putative | N/A |
| PBANKA_1437100 | conserved Plasmodium protein, unknown function | N/A |
| PBANKA_1441700 | parasitophorous vacuolar protein 2 | PV2 |

**Table S3.** Top most upregulated genes at 12 hours post-infection.

| Gene ID | Product Description | Gene Name |
|---|---|---|
| PBANKA_0108900 | peptidyl-tRNA hydrolase PTRHD1, putative | N/A |
| PBANKA_0201000 | fam-b protein | N/A |
| PBANKA_0201500 | Plasmodium exported protein, unknown function | N/A |
| PBANKA_0211500 | proteasome subunit alpha type-5, putative | N/A |
| PBANKA_0315900 | pseudouridine synthase, putative | N/A |
| PBANKA_0404200 | ubiquitin-conjugating enzyme E2, putative | N/A |
| PBANKA_0406500 | T-complex protein 1 subunit eta, putative | CCT7 |
| PBANKA_0511900 | 60S ribosomal protein L3, putative | RPL3 |
| PBANKA_0517200 | haloacid dehalogenase-like hydrolase, putative | HAD1 |
| PBANKA_0619100 | 40S ribosomal protein S5, putative | N/A |
| PBANKA_0619300 | conserved Plasmodium protein, unknown function | N/A |
| PBANKA_0623200 | lysophospholipase, putative | N/A |
| PBANKA_0623500 | fam-a protein | N/A |
| PBANKA_0700500 | fam-a protein | N/A |
| PBANKA_0922800 | small nuclear ribonucleoprotein Sm D1, putative | SNRPD1 |
| PBANKA_1011900 | H/ACA ribonucleoprotein complex subunit 3, putative | NOP10 |
| PBANKA_1101100 | Plasmodium exported protein, unknown function | N/A |
| PBANKA_1101200 | Plasmodium exported protein, unknown function | N/A |
| PBANKA_1120200 | pyridoxine biosynthesis protein PDX1, putative | PDX1 |
| PBANKA_1301400 | proteasome subunit alpha type-1, putative | N/A |
| PBANKA_1302241 | tRNA Selenocysteine | N/A |
| PBANKA_1305100 | 60S ribosomal protein L1, putative | N/A |
| PBANKA_1326400 | glyceraldehyde-3-phosphate dehydrogenase | GAPDH |
| PBANKA_1360300 | DNA/RNA-binding protein Alba 4, putative | ALBA4 |
| PBANKA_1365680 | fam-c protein | N/A |
| PBANKA_1403000 | small heat shock protein, putative | N/A |
| PBANKA_1407600 | 60S ribosomal protein L24, putative | N/A |
| PBANKA_1446200 | mitosis protein dim1, putative | N/A |

### Data Files

**Data S1. (xlsx). Differential expression**. Gene expression (RNA-seq) of *P. berghei* for all the datasets analyzed in this study. Fold change was determined for each time point vs the sporozoite samples. Columns are the following: GeneID; GeneName: gene product description (PlasmoDB); IsCoding: is the gene a known protein-coding gene?; LogFC: Log2(fold-change); lfcSE: standard error of the log2(fold-change); stat: Wald test statistic; pvalue; padj: FDR-corrected p-value; <SampleID> Normalized expression value for specific <SampleID>.

**Data S2. (xlsx). Cluster assignment**. This file shows the genes that belong to each cluster. Columns are the following: GeneID; GeneName: gene product description (PlasmoDB); Cluster: cluster number.

**Data S3. (xlsx). Cluster analysis**. Characterization of the 14 co-expression clusters. Columns are the following: Cluster: cluster number; GO terms: GO terms associated with each cluster ($p < 0.01$); Notable genes: selected genes from each cluster; DNA motif: enriched DNA motifs for each cluster discovered through the DREME (e-value < 0.05); E-value: enrichment value for each DNA motif.
